# Role of competition in the strain structure of rotavirus under invasion and reassortment

**DOI:** 10.1101/137661

**Authors:** Daniel Zinder, Maria A Riolo, Robert J. Woods, Mercedes Pascual

## Abstract

The role of competitive interactions in the formation and coexistence of viral strains remains unresolved. Neglected aspects of existing strain theory are that viral pathogens are repeatedly introduced from animal sources and readily exchange their genes. The combined effect of introduction and reassortment opposes strain structure, in particular the predicted stable coexistence of antigenically differentiated strains under strong frequency-dependent selection mediated by cross-immunity. Here we use a stochastic model motivated by rotavirus, the most common cause of childhood diarrheal mortality, to investigate serotype structure under these conditions. We describe a regime in which the transient coexistence of distinct strains emerges despite only weak cross-immunity, but is disturbed by invasions of new antigenic segments that reassort into existing backgrounds. We find support for this behavior in global rotavirus sequence data and present evidence for the displacement of new strains towards open antigenic niches. Our work extends previous work to bacterial and viral pathogens that share these rotavirus-like characteristics, with important implications for the effects of interventions such as vaccination on strain composition, and for the understanding of the factors promoting emergence of new subtypes.

## INTRODUCTION

Understanding how pathogen variation is generated, structured and maintained is a central question in disease ecology and is of fundamental importance to epidemiology. Ultimately, such research requires the synthesis of a wide set of fields including evolutionary, epidemiological, immunological, and ecological dynamics^1^. A major body of theoretical work at this intersection posits that the combined immunity of the host population can drive pathogens to differentiate into groups with reduced immune cross-reactivity, as intermediate hybrids are suppressed by the protection of the population to the established parental strains^2–7^. This organization into discrete strains as the result of ‘immune selection’ relates to ideas at the core of theoretical ecology including competitive exclusion, niche differentiation, character displacement and limiting similarity, suggesting that species cannot stably coexist when using a single resource, and that the boundaries of realized (or occupied) niches are set by competition^8–12^. However, in previous models of niche differentiation the dynamic role of evolution in the maintenance and formation of these niches was only partially explored, and establishing empirical evidence for a role of non-neutral processes in the coexistence of diverse communities has been challenging^12–14^.

In the context of pathogens, the role of mutation and recombination in niche differentiation has been evaluated under competitive interactions mediated by a high level of serotype-specific immunity and for a restricted set of antigenic variants from a ‘finite pool’^4–6^. Here, instead we consider low levels of specific immunity and an unlimited antigenic reservoir so that we can address the structure and dynamics of serotype diversity in rotavirus A (RVA), the leading cause of diarrheal deaths in children worldwide. RVA experiences recurrent introductions of antigenic novelty through zoonotic transmission, as well as frequent reassortment between its segments and weaker specific (homotypic) than generalized (heterotypic) immunity. These are additional processes which potentially undermine strain structure and coexistence. For pathogen populations, antigenic variants are the traits mediating frequency-dependent competition for hosts, with an advantage of the rare and the disadvantage of the common, akin to stabilizing competition between species^15^. We ask whether strain structure can still emerge from competition for host immune space under these rotavirus-like conditions, and whether there are signatures of such non-neutral process in global sequence data.

Rotavirus A (RVA) is a non-enveloped virus, with 11 double stranded RNA genome segments. Two outer capsid proteins VP4 (P types) and VP7 (G types) are the major antigens, often used for classification of rotavirus into genotypes (e.g. G1P[8], G3P[8], G9P[6]) because of their role in the generation of humoral immunity^16,17^. There is evidence of increased protection against infections by the same genotype ^18,19^ yet a large component of immunity is heterotypic^20–22^. At a segment level, each segment type (e.g. G1, G12 etc.) corresponds to a monophyletic cluster, and includes either single or multiple cross-species transmission into humans from animal sources^23^. Although mutation is responsible for a certain degree of antigenic change within individual G types (e.g. within G1), its role in the antigenic evolution of rotavirus within the human population is thought to be limited^24,25^. Multiple G and P types have been described, and in contrast to influenza, serotypes coexist and undergo frequent reassortment^26,27^. Several of the discovered G and P types are more abundant than others, and only a small fraction of all possible genotypes of RVA have been reported^17,20,26,28^. This has been attributed to a balance between preferred genome constellations and reassortment^26^, and there is evidence that these constellations have an improved fitness unrelated to immunity^29^. However, it should be noted that in the absence of specific immunity or an alternative type of niche differentiation (e.g. host and tissue tropism), hosts act as a single resource and simple ecological and epidemiological models predict competitive exclusion by the strain with the largest basic reproductive number^8,30^. Mechanisms for stable co-existence in the absence of this type of niche differentiation exist but have not been observed for rotavirus^31–34^.

The frequency of circulating rotavirus serotypes varies over time and across global geographic regions^28^. On a local level circulating serotypes are often partially or fully replaced after being common in a region for several years^35–37^, yet those can re-emerge at later periods. These replacement dynamics, taken together with possible changes in serotype prevalence following vaccination^38–40^ suggest that selective pressures generated by host immunity can drive changes in circulating serotypes. However, the role of such immune selection (stabilizing competition) in the maintenance and structure of rotavirus serotypes has not been established. Motivated by important challenges in rotavirus, we formulate a stochastic transmission model including antigenic evolution to investigate how serotype structure is generated and maintained. Our work extends the dynamic regimes previously identified by strain theory and phylodynamic models, to include the introduction of antigenically novel segments from the zoonotic introduction of alleles^1,2,5,6^. Empirical patterns in the global sequence data for the virus are consistent with our simulation. An improved model of how strain communities are generated and maintained provides a basis to also understand their expected response to human interventions, as well as the factors that promote or hinder the invasion of new subtypes and serotypes in rotavirus and other pathogens.

## RESULTS

### Strain structure in a model motivated by rotavirus

An individual-based SIR model explicitly tracks the chains of infection by viral lineages as well as the antigenic phenotype of every virus in the population. A virus in the model is composed of three antigenic segments (e.g. A, B, C), each segment allele (e.g. A_3_) representing a unique serotype, with a combination of such alleles representing a viral strain (e.g. A_1_B_2_C_2_). Novel alleles (e.g. C_3_) are introduced into existing backgrounds (e.g. A_1_B_2_C_2_ → A_1_B_2_C_3_). The model was modified from previous models^6,41^ to include reassortment and the genealogical tracking of each viral segment independently. Immunity and infectivity are also different from the cited models, and in particular both generalized (heterotypic) and specific (homotypic) immunity are included, as described in detail in Materials and Methods.

One striking feature of rotavirus epidemiology is the dramatic shift in serotypes in the absence of major shifts in yearly incidence. This pattern contrasts with that of other viruses such as influenza where the emergence of new antigenic variants typically implies epidemic behavior with an increase in attack rates^42^. To capture the year-to-year variation in incidence, we calculated the two-year coefficient of variation in rotavirus positive cases from 2001-2012 from a cohort study of infants in Bangladesh^36^, as CV=0.04, where CV is the standard deviation divided by the mean fraction of rotavirus positive cases. To compare how variable, or stable, infection levels in our simulations are, we plot the coefficient of variation for prevalence with varying parameters.

Parameter ranges for generalized and specific immunity were chosen to match published rotavirus clinical data. The range of generalized immunity considered, was determined by fitting the probability of reinfection in a cohort of Indian children (σ_gen_=0.4) and including a wide range above and below that estimate (σ_gen_=0.2-0.6)^43^ (**Figure S1**). With this estimate of generalized immunity, we observe stable infection levels, characteristic of rotavirus when there is low specific immunity (σ_spec_=0.25) (**Figure S2**). As the value of specific immunity is increased, epidemic dynamics coupled by the full or partial sequential replacement of strains occur, similar to influenza, (**Figure 1A**).

**Figure 1.**
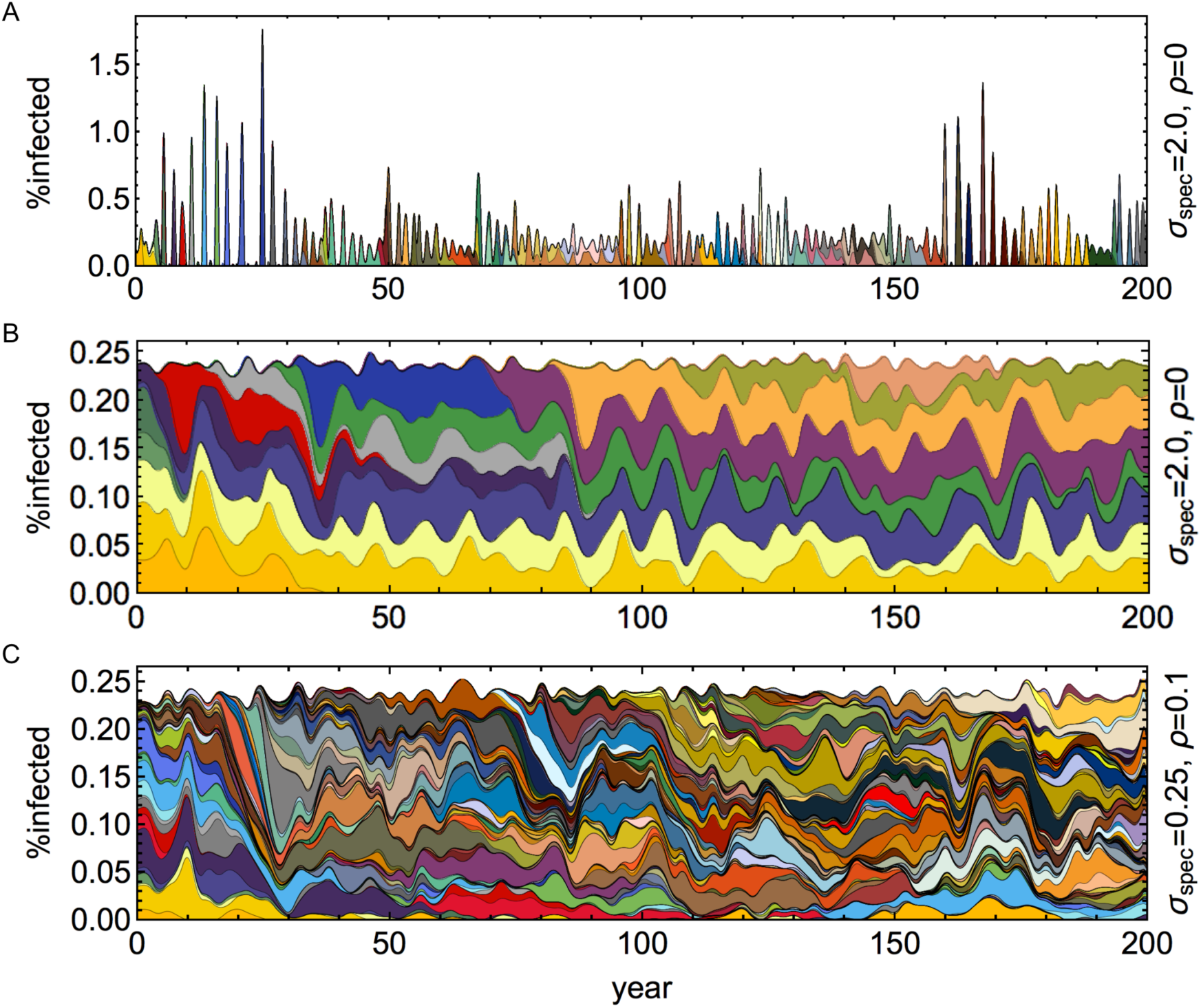
Contrasting serotype dynamics with varying specific immunity and reassortment rates. Cumulative prevalence, color coded by serotype (unique combination of three segments) for varying specific immunity (σ_spec_) and reassortment (ρ) rates. The specific example in **A** illustrates the replacement regime. **B** and **C** illustrate the coexistence regime with reassortment. **A.** High specific immunity (σ_spec_ = 2.0), leading to the sequential replacement of serotypes coupled with high incidence variability and epidemic dynamics. **B.** With low specific immunity σ_spec_ = 0.25 in the absence of reassortment (ρ = 0) invasions occurring on existing backgrounds have a temporary frequency-dependent advantage and gradually displace existing serotypes. **C**. In the presence of reassortment (ρ = 0.1), invading segments reassort with multiple backgrounds and transiently disrupt serotype structure. (additional parameters: β=3.51, σ_gen_=0.4, i_I_=8 year^-1^)

We then moved to characterizing the role of reassortment in the setting of rotavirus-like epidemiology and immune modeling. Reassortment rates had little effect on prevalence variability in the simulations when coupled with low levels of specific immunity characteristic of stable levels of infection, as assessed by coefficient of variation over two-year time windows (**Figure S2**). However, reassortment had a major impact on the genetic structure of the population (**Figure 1B, C)**. This genetic structure is measured in two ways. First, Simpson diversity, representing the effective number of segment types at a locus, and can be calculated as one over the probability that two serotypes randomly selected from a sample will share the same segment at the locus. Second, we measure the amount of niche overlap as the fraction of shared segment alleles between strains from randomly sampled infected hosts, excluding antigenically identical strains. In the absence of serotype-specific immunity (σ_spec_=0), segment alleles are neutral, and a single serotype persists for the duration of the simulation (**Figure 2B**). In previous studies, in the context of a finite antigenic pool and strong specific immunity, reassortment had a limited effect on strain structure^4^. Here, we see that with lower levels of specific immunity and with the repeated introduction of antigenic novelty, reassortment results in increased niche overlap, higher diversity of strains, and more complex strain dynamics (**Figure 1B, C**, **Figure 2A, C, D, E**). As expected, however, the formation of a strain structure with reduced niche overlap is substantially diminished with higher reassortment rates (**Figure 2D, E**).

**Figure 2.**
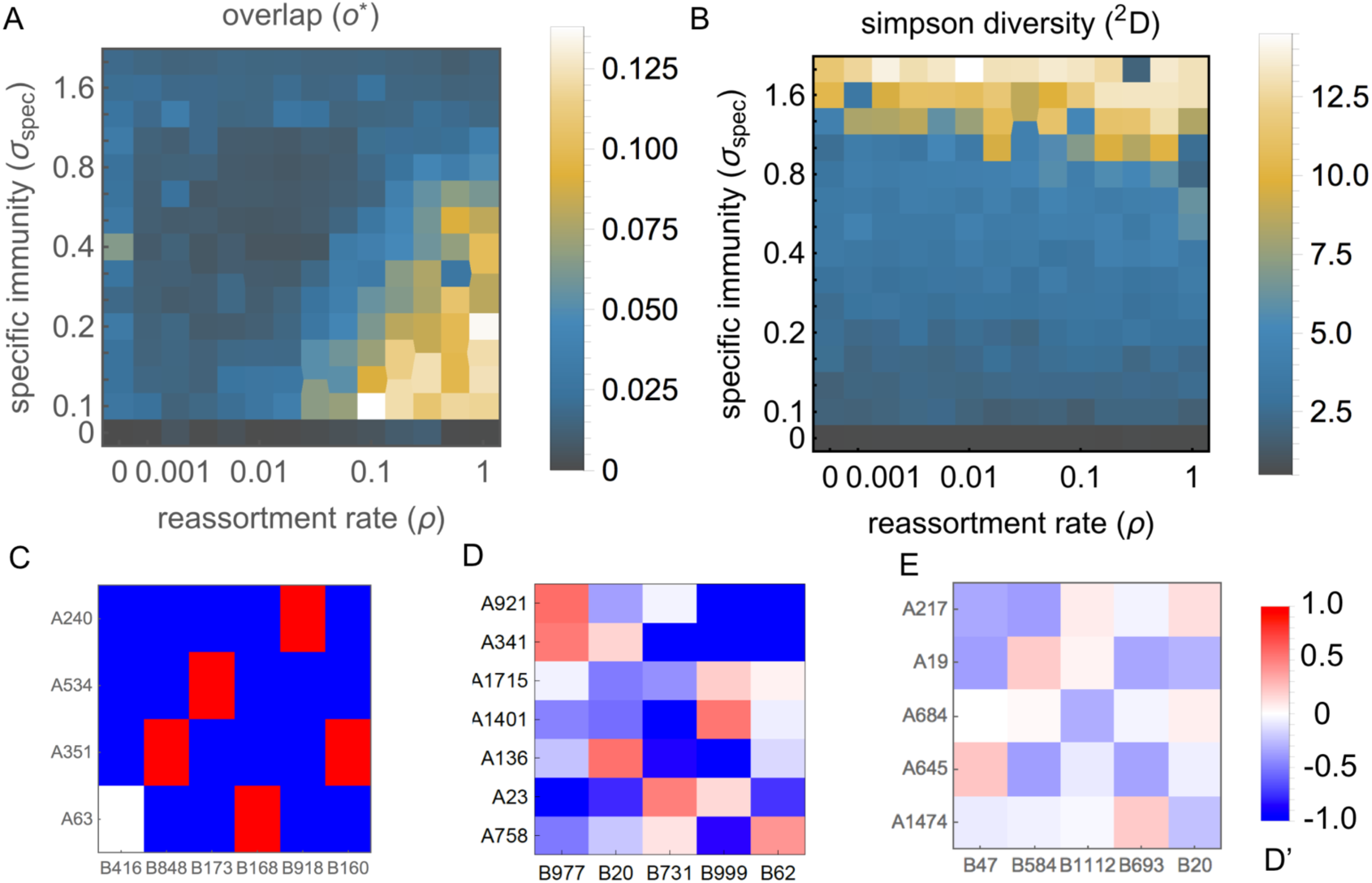
Serotype community structure with varying specific immunity and reassortment rates. **A.** The mean segment overlap between serotypes as a function of specific immunity and the rate of reassortment. Segment overlap was calculated as the fraction of shared segments between randomly sampled (non-identical) serotypes during 200 simulated years (parameters: β = 3.51, σ_gen_ = 0.4, t_I_=8 year^-1^). **B.** The Simpson diversity (1/λ or ^2^D) of one of three antigenic loci during 200 simulated years, with varying specific immunity and reassortment rates. In the absence of specific immunity, a single segment serotype is present determined by the random fixation of a newly introduced alleles. With increasing specific immunity, segment diversity increases with their fixation promoted by a frequency-dependent competitive advantage. Reassortment promotes segment diversity for intermediate levels of specific immunity. **C.** The mean normalized linkage disequilibrium (D’) between two segments at the first two of three antigenic loci. Without reassortment, clear linkage appears only in combination with a single other segment, indicating the reduced prevalence of strains which share more than one segment. (σ_spec_ = 0.25, ρ = 0). **D.** Linkage disequilibrium with increasing reassortment rate (σ_spec_ = 0.25, ρ = 0.1) **E.** Increasing overlap and reduced linkage disequilibrium with an even higher reassortment rate (σ_spec_ = 0.25, ρ = 1).

In previous models which generate discrete strain structure, antigenically differentiated strains were maintained through the suppression of recombinant strains by the combined immunity of the host population against the parental strains^2^. Here, additional competitive forces exist in the form of selection which favors strains carrying novel antigenic segment alleles, specifically in relation to their parental backgrounds; evidence for which is seen in the form of largely non-overlapping strain structure indicating the extinction of the background strains on which new segment alleles were introduced (**Figure 2C**). Importantly, with reassortment new segments can travel across multiple backgrounds. Following an introduction, reassortment is initially favored; allowing the introduction of new invading alleles which quickly associate with multiple backgrounds until they reach their frequency dependent equilibrium and the benefit of reassortment diminishes (**Figure S3A**). This movement of new segments through multiple backgrounds transiently destabilize the otherwise stable coexistence of largely non-overlapping strains (**Figure 1B-C**).

Surprisingly, at low specific immunity levels, this change in the composition of strains (**Figure 1B, C**) with increasing reassortment rate has a limited effect on the prevalence and diversity of individual segments (**Figure 3A, B**). The genealogy resulting from these simulations is comparable to the inferred genealogy of rotavirus (**Figure S4**) in the presence of long term co-existence of distinct segment genotypes. A degree of stability in the prevalence of established types, despite changes in strain composition is consistent with observed patterns in epidemiological surveys of rotavirus^36^.

**Figure 3 A-B.**
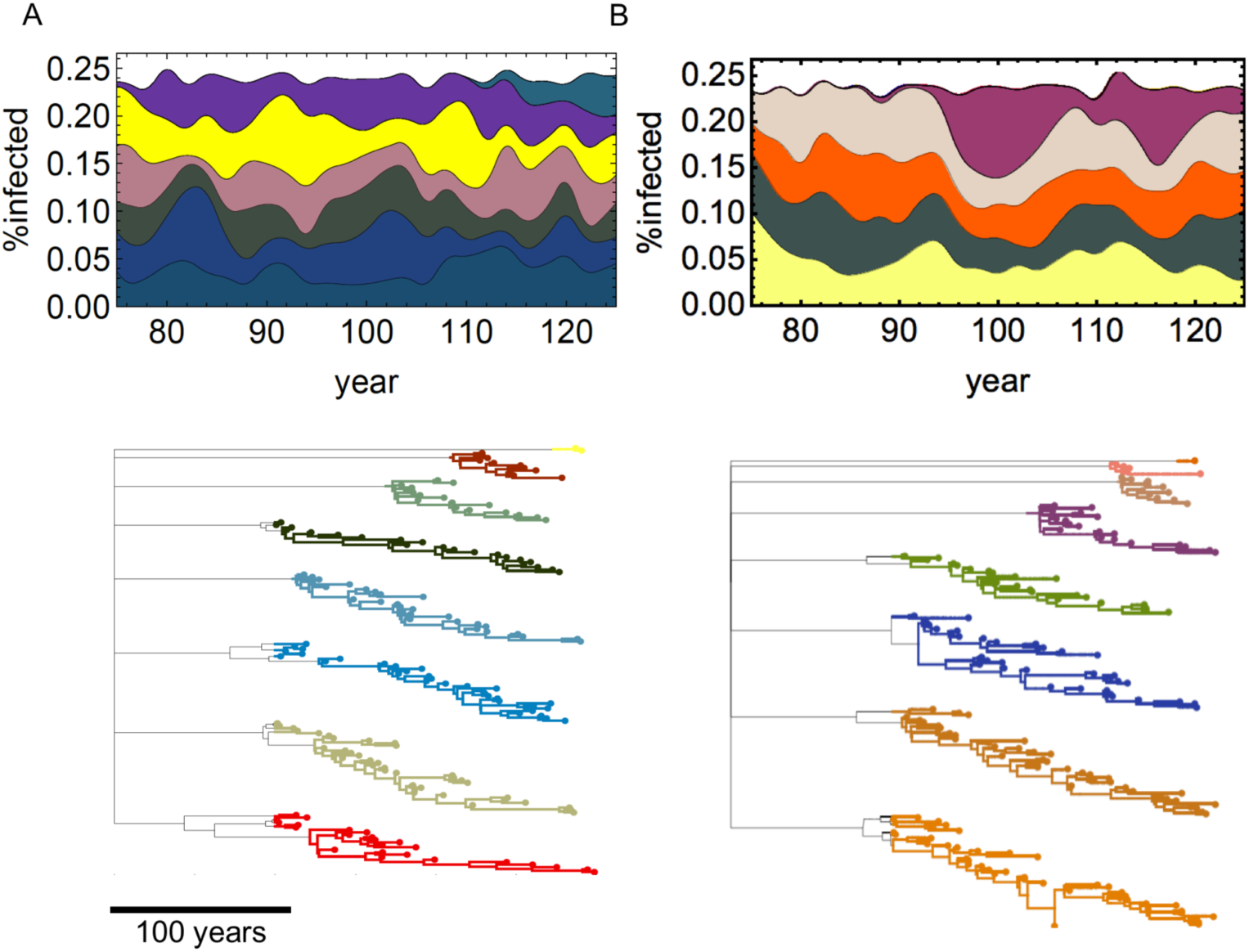
Cumulative prevalence and simulated genealogy. Cumulative prevalence now at a single locus, color coded for the different segment serotypes. The corresponding simulated genealogy is also shown. New segment serotypes are simulated to share an ancient source of coalescence in a source of unlimited diversity. **A**. low reassort rate σ_spec_ = 0.25, ρ = 0.1 **B**. high reassortment rate σ_spec_ = 0.25, ρ = 1. Changes in the relative prevalence of individual segments are less pronounced in comparison to changes in strain composition (**Figure 1**). The simulated genealogy is comparable to the genealogy of rotavirus G protein (**Figure S4**) in the presence of long term co-existence of distinct segment genotypes.

### Strain structure from empirical rotavirus sequence data

To measure the number of associations of different G and P types in rotavirus we use a sample of globally available *rotavirus A* genotypes collected from publically available sequence data between 1990-2015^27^. Before 1995 four G genotypes (G1, G2, G3, G4) were the most prevalent worldwide in humans, and typical G-P combinations including G1P[8], G2P[4], G3P[8] and G4P[8] were responsible for the majority of rotavirus diarrhea among children^28^. An additional genotype G8, of likely bovine origin has been identified as a common cause of rotavirus infection in humans, especially in Africa during the 1990s^44,45^. Since 1995 the G9 genotype has become increasingly more common and has been observed in association with multiple P type backgrounds^23,46,47^. In addition, G12 genotype strains have been found to circulate in most parts of the world since the start of the new millennium^23,48^. In both the case of G9 and G12 a single lineage was found to be responsible for the majority of worldwide spread with a suspected origin in pigs, while multiple previous human infections have been recorded^17^.

We compare the diversity of G-P genotypes, contrasting between established (G1, G2, G3, G4) and more recently introduced (G6, G8, G9, G12) G-types as defined in^26,28,49^. In **Figure 4A** we plot the sampling frequency of common rotavirus genotypes grouped by sharing a G protein genotype/serotype^50^. G types that have become more prevalent recently show associations with multiple P types, consistent with the model’s behavior of movement through different established backgrounds. By contrast, the established genotypes are associated with fewer P backgrounds (p=0.03, T-test two-tailed): thus, G1-G4 are associated on average with 1.2 P-types (Inverse Simpson index) compared to 2.3 types for the more recent G6, G8, G9 and G12 (**Figure 4B**). Similarly, P6, which has recently become more widespread is associated with 5.9 G-types compared to 1.8 G-types on average for P4 and P8 (**Figure 4C**) (p=0.01, likelihood ratio test comparing a single normal-distribution with two sharing a common variance). The decrease diversity of associated backgrounds for established types is not observed in the model, a departure we address in the discussion.

**Figure 4.**
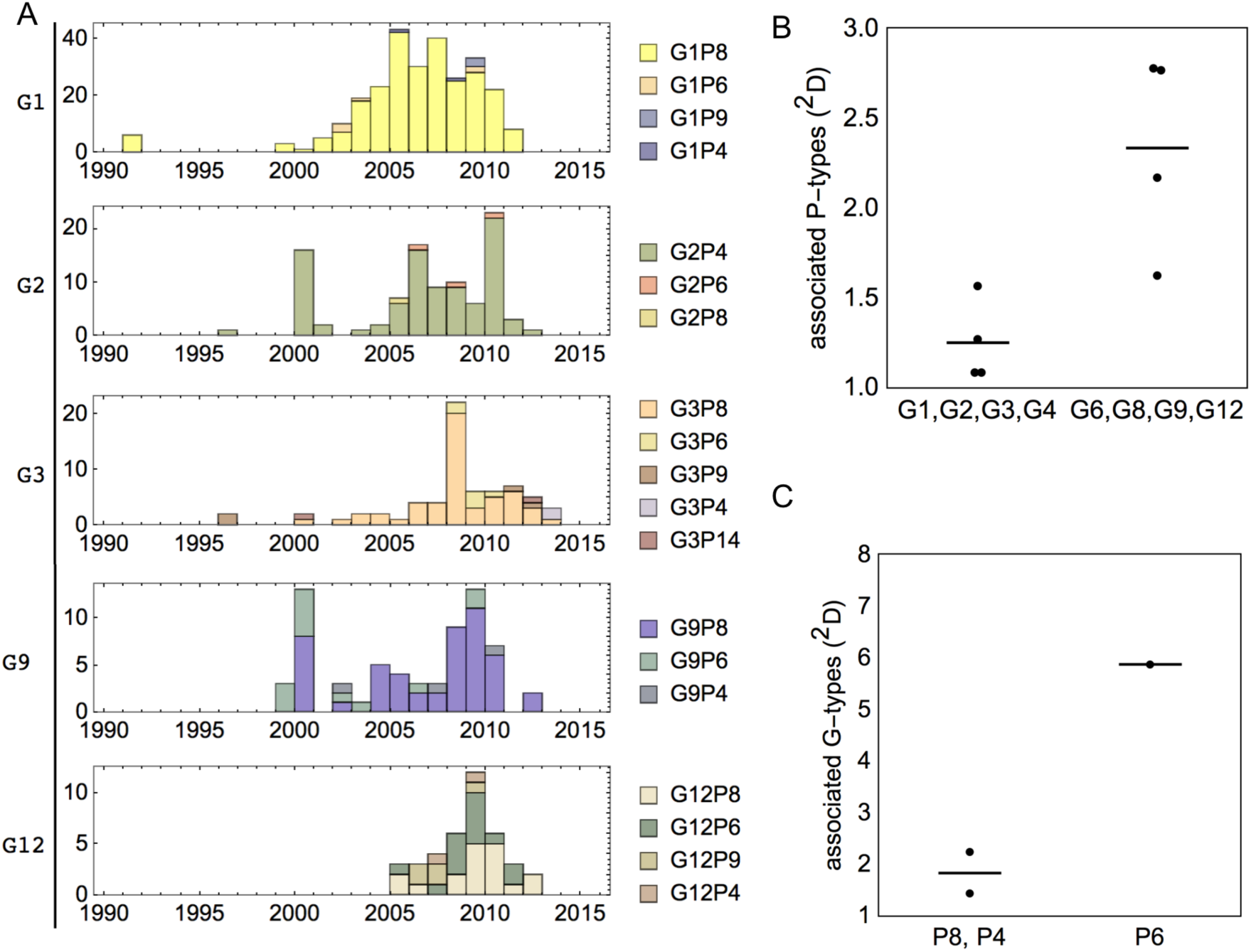
Diversity of association in publically available G-P genotypes. a sample of 588 globally available human rotavirus A genotypes collected from publically available sequence data between 1990-2015 (Woods et al. 2015) **A.** Cumulative sample timeseries for G-P genotypes grouped by their G type, showing representative ‘established’ G types (G1-G3) and more recently widespread G types (G9-G12) **B.** Simpson diversity of publically available G-P genotypes contrasting between established (G1-G4) and more recently widespread (G6, G8, G9, G12) G-types. **C.** Simpson diversity of publically available G-P genotypes contrasting between established (P8, P4) and more recently widespread (P6) P-types

Next, we examine empirical evidence of strain structure in the form of weakly overlapping niches as predicted by theory. We plot the abundance of different G-P genotype combinations (**Table S1**) in matrix form, ordered from the most prevalent to the least prevalent segment types (**Figure 5A**). In addition, we plot the p-value for a specific G-P pair being above or below its expected abundance if segment associations were random while maintaining the same number of segments for each type. We observe significant departures from random expectations based on frequency. These departures show a checkerboard pattern of enhanced and reduced linkage, indicating reduced strain overlap (o=0.06, p<0.00001, N=668) in comparison to the random expectations based on and suggestive of niche differentiation through competitive interactions. In the simulation, such patterns of linkage were observed in the presence of specific immunity and with a limited amount of reassortment. In the absence of specific immunity, competition for a single resource prevented the co-existence of multiple strains, while high levels of reassortment resulted in reduced niche differentiation and a smaller deviation from random linkage expectations.

**Figure 5.**
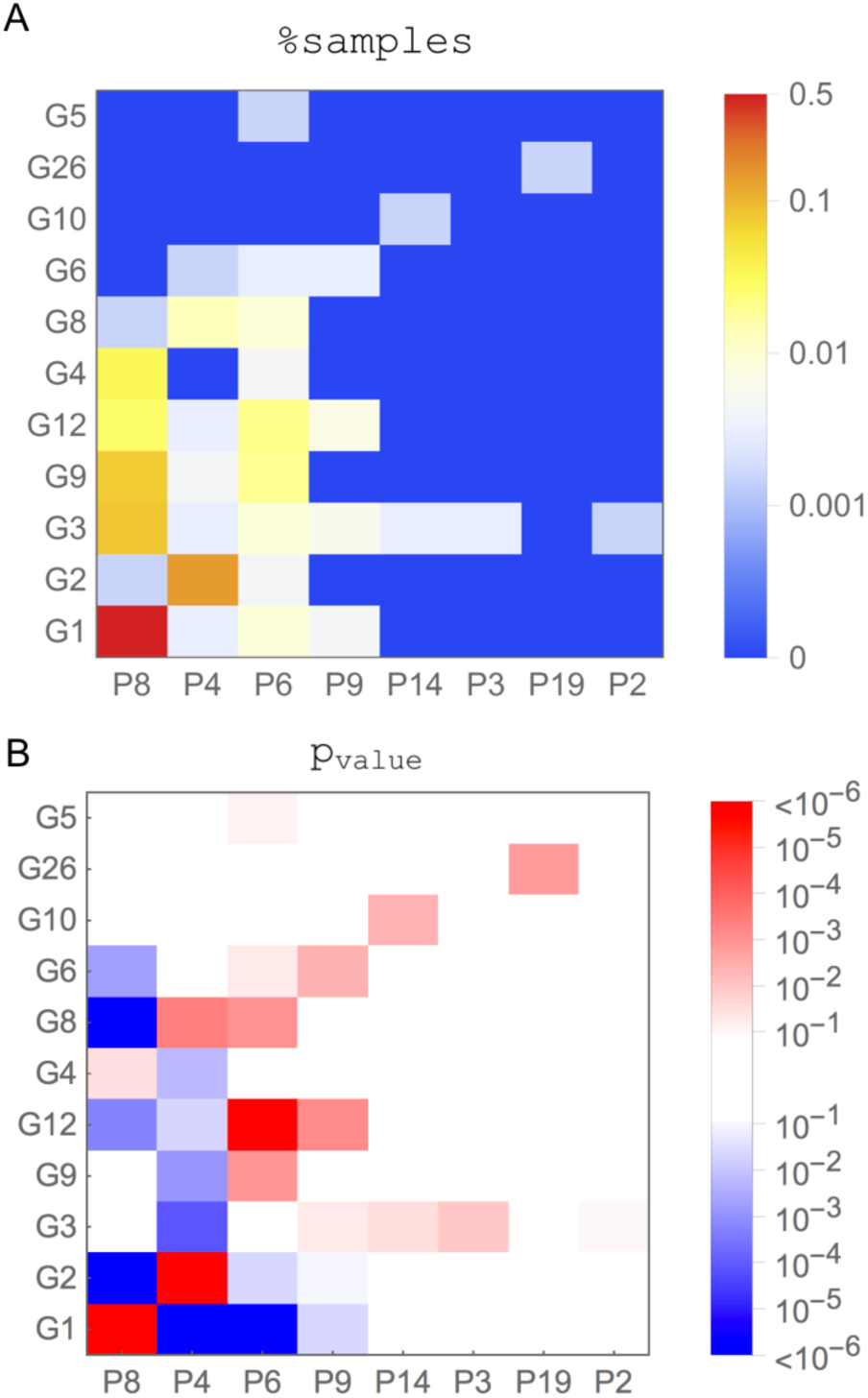
Sampling and deviation from equilibrium frequency for publically available G-P genotypes. **A** The percent of samples for different G-P genotype combinations in publically available sequence data^27^. P and G segments are ordered by their prevalence. **B** The p-value for a specific G-P pair being above or below it’s expected frequency. P-values were calculated using Fisher’s exact test between G-P pairs using a two-tailed test and are colored red or blue based on the pair abundance being below or above its random expectation (respectively). Deviations from expected frequencies establish a checker-board pattern, consistent with selection for low overlap between dominant strains (e.g. G1P[8] and G2P[6]), seen in the lower-left corner. Interestingly, positive deviations for the associations of more recently invading segments (e.g. G12) are with P types that are not found in frequent partnerships already, such as P6. Note the corresponding dominance of red boxes towards the right and upper corner, which represents a part of immunity space in the host population that should be more open, given the low frequency of the corresponding pairs and segments.

The most common and non-overlapping combinations of G-P types, G1P[8] and G2P[4], have a higher abundance than expected based on segment prevalence alone (**Figure 4B**). G3 ranking next in abundance is associated with multiple backgrounds, with the association with P[6], P[9] and with P[3] in a higher proportion than expected by chance, conforming to non-overlapping combinations with the most common genotypes G1P[8] and G2P[4]. For the more recent G-types, G9 and G12 ranking next in their abundance. Despite having high abundance in association with multiple backgrounds, in particular with P8 which can be trivially expected from the high frequency of this P type, these more recent G-types are more abundant than expected in association with rarer P types including P6 and P9. These positive deviations are therefore seen for associations that would constitute relatively open niches in the immune landscape of the host. Similarly, negative deviations, for example for G12P[8], occur for associations with segments such as P8 that are already common in resident strains. The inferred origin of recent G type invasions to humans (such as G9 and G12) from a single lineage^17^, suggests that the observed associations with multiple backgrounds are the consequence of post invasion reassortment which is not limited to the initial invading background. Furthermore, these multiple associations are consistent with invading G-types and P-types favoring association with rarer backgrounds as the result of the limited availability of open niches. In addition, rarer G-types (G6, G10, G26) have higher abundance than expected by chance in association with less common P types (P9, P14, P19). Although it is likely that these associations represent the original association present during spillover events, we should consider the possibility that such spillovers would not have occurred in the context of occupied niches.

Rotavirus displays temporal and spatial variation in strain composition. To address patterns of niche differentiation at this scale we interrogate sequence data locally, at the country of origin level, and based on decade long time intervals (1985-1995, 1995-2005, 2005-2015). We calculated the community overlap of genotypes in a specific country and decade and the probability that this or a lower overlap of was generated by chance in a random assembly of global segment alleles. We find that e.g. in 2005-2015, for several countries including e.g. the U.S. (o=0.16, p=0.013, n=54), China (o=0.05, p<0.0001, n=27), Australia (o=0.012, p<0.0001, n=85) and Paraguay (o=0.16, p=0.008, n=33), serotype overlap was significantly lower in comparison to random expectation (**Figure S5**, **Figure S6**), a pattern consistent with immune mediated niche partitioning. In addition, the associations that are significantly higher than expected at random, often vary between different countries supporting the role of immune mediated niche partitioning.

## DISCUSSION

The role of competitive interactions in the formation and maintenance of viral strains is a subject of ongoing investigation. This research complements the emphasis that has been given to interspecies transmission events in the emergence of viral subtypes and serotypes, and to epistatic interactions limiting the associations between different segment alleles. In this study, we evaluated the role of immune-mediated competitive interactions in the formation of serotypes in a model that captures major aspects of rotavirus dynamics, and is applicable to other viruses, in particular ones which encounter reassortment in the face of the repeated introduction of antigenic novelty from a diverse source, such as the zoonotic reservoir, and ones with limited type-specific immunity. We show that under these conditions strain structure can emerge, and involves a balance between competitive forces driving immune-mediated niche differentiation, and disturbance to this structure by repeated invasions and frequent reassortment. Existing rotavirus sequence data exhibits a non-random structure which is consistent with this dynamical regime.

In contrast to viral infections such as influenza, where antigenic changes have been shown to result in increased epidemic size, serotype turnover in rotavirus has limited impact on yearly incidence^36,51^. We capture this type of behavior in our simulation when specific immunity is sufficiently low. The presence of specific immunity, even at relatively modest levels, confers a selective advantage to newly introduced segments, and promotes their invasion. The selective advantage of invading segments while they are rare promotes their association with multiple backgrounds through reassortment. Segments invading on existing backgrounds, can lead to the extinction of their background strains. For low levels of reassortment a degree of niche differentiation between strains is maintained. However, there is a limit to the ability to maintain strain structure and with low enough specific immunity, reassortment can readily break the signature of full niche partitioning.

Evolutionary and ecological processes other than immunity take part in generating pathogen strains. For instance, different subtypes of influenza A (i.e A/H1N1, A/H3N2, A/H5N7) represent separate spillover events from an avian reservoir, with a possible intermediate host^52^. While reduced cross-immunity in this case appears as a by-product of non-immune processes, and strong epistatic interactions are present^53^, mechanisms similar to the ones suggested in the model are evident in the maintenance of strain structure in influenza. Reassortment between subtypes combined with competitive interactions has allowed the displacement of one influenza subtype A/H2N2 by another A/H3N2^54^, and antigenic shift allows for novel viral antigens which are less frequent, such as pandemic A/H1N1 to invade^55^. It is probable that immune mediated frequency dependent competition may also be at play in allowing spillovers from specific subtypes such as A/H5N7 to occur, while preventing e.g. the reintroduction of existing subtypes from occupied niches^56^. Additional non-immunity based processes, such as geographical isolation, tissue, and host tropism can also generate viral strains^57–59^. Importantly, non-immune mechanisms can work in tandem with immune mediated ones to generate and maintain pathogen strains and to reinforce niche differentiation. Both are likely to play a role in shaping diversity patterns in rotavirus.

Consistent with the simulation, we observe the rapid association of recently introduced G and P types with diverse partners (**Figure 4**). However, in contrast with simulation, established serotypes maintain a fixed number of associations for prolonged durations (**Figure S3A**). We attribute the higher number of associations of older G and P types to missing components in our model, namely, a higher fitness for preferred genome constellations^26^. When a ‘crystallization’ mechanism was introduced, overall association diversity was reduced, and when reassortment was sufficient to generate multiple associations newly introduced segments showed an increasing diversity of association followed by a decline (**Figure S3B**). Further, research into this suggested mechanism and the role of fitness differences, in generating this and additional patterns of strain diversity in rotavirus is necessary. An additional missing component is the composition of additional genome segments, and their organization into genogroups. Instead, to maintain the simplicity of our model we considered three equivalent antigenic segments. Finally, a model which considers population sizes, and effective population sizes and structure, consistent with the ones observed in rotavirus may help in future research and in distinguishing neutral from non-neutral evolutionary patterns. However, rotavirus strains experience global circulation and there is no clear association between different strains and geographic regions, suggesting geographic population structure isn’t sufficient to maintain the observed serotype dynamics.

We have shown that a coexistence regime exhibits in rotavirus a non-random structure with a signature of competitive interactions and weakly-overlapping niches between strains. Our results suggest that dramatic changes in serotype composition can occur without a dramatic change in standing generalized immunity. Moreover, targeted interventions, identifying shared commonly recognized antigens, may identify transient sweeping alleles as being most common. These alleles may have no inherent fitness differences compared to other alleles, with the exception of a transiently higher host susceptibility. As such, targeted vaccination against these alleles may not offer higher efficacy. Finally, we expect some of the barriers on the formation and on the invasion of strains to originate from immune mediated competition, which can change in direction and composition with the introduction of vaccination, potentially leading to unexpected changes to strain composition. Reassortment, combined with zoonosis has been the mechanism behind the emergence of major recent pandemics such as the 2009 H1N1 influenza pandemic. It is therefore crucial to understand the role the population susceptibility, specific and generalized, in preventing such spillover events from animal sources, and to understand the potential of viral segments to reassort. As is the case with influenza, the study of the global diversity of rotavirus strains, both human and zoonotic is important in preventing the emergence of invading serotypes. All else being equal, a polyvalent vaccine leading to a higher vaccine breadth, would help reduce the potential for new strains which are substantially different from the vaccine strain to invade or form through reassortment. A dramatic shift in serotype composition may result in small but important differences in immunity; a reduction in efficacy of 5-10% can nonetheless result in tens of thousands of lives.

## METHODS

**Table 1.** Model Parameters

| description | symbol | value | ref |
| --- | --- | --- | --- |
| contact rate of symptomatic infection | $\beta$ | $1.0 \text{ day}^{-1}$ or $3.514 \text{ day}^{-1}$ | <sup>60</sup> |
| duration of infection | $\frac{1}{\rho}$ | $7.0 \text{ days}$ | <sup>60</sup> |
| homotypic immunity | $\sigma_{gen}$ | $0.2-0.6$ | <sup>43,60,61</sup> * |
| heterotypic immunity | $\sigma_{spec}$ | $0-2.0$ | |
| mutation rate | $\mu$ | $0$ | |
| introduction rate | $i_I$ | $0-32 \text{ year}^{-1}$ | |
| probability of first infection to be symptomatic | $x_0$ | $0.47$ | <sup>60</sup> |
| Reduction in infectivity with subsequent infections | $\xi$ | $0.62$ | <sup>60</sup> |
| Population size | $N$ | $10,000,000$ | |
| number of viral segment types | $N_S$ | $3$ | |

\* also see **Figure S1**

### Model Description

In previous models, strong immunity against each segment allele, was necessary for antigenic niche differentiation between strains ^2,4,40^. Cohort studies and vaccine studies suggest rotavirus departs significantly from these assumptions in that: 1. multiple infections are possible, with a diminishing probability based on the number of previous infections, and 2. much of the immunity gained is generalized, that is a host infected with one strain gains some portion of protection against all strains 3. the portion of immunity that is specific to a strain is debated ^20–22^. To better match these aspects of rotavirus epidemiology, we explore a range of parameters and explicitly include both a generalized and a strain specific component to immunity. In our model, the risk of infection has a generalized component which follows an exponential decline based on the number of previous infections with any rotavirus strain and a specific component which follows an exponential decline based on the fraction of segments in the current strain which have been seen before by the host in previous infections. In contrast with previous models, we include the arrival of newly introduced segment alleles ^36^. These are introduced in the simulation at a given annual rate. Reassortment is also modeled, and when a host is infected with multiple viral strains segments are reassorted with a certain probability. Model parameters are listed in **Table 1**.

More specifically, a host’s risk of infection follows an exponential decline based on the number of infections and the number of unique antigenic segments to which that the host has previously been exposed:

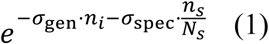

where *σ_gen_* is the level of heterotypic immunity, *σ*_*spec*_ is the level of homeotypic immunity, *n*_*S*_ is the number of infections a host has already experienced, *n*_#_ is the number of segments the host has been exposed to previously, and *N*_*S*_ is the number of antigenic segments of the virus has (*N*_*S*_=3 for all the simulations reported here). Immunity is gained upon recovery from an infection.

Only symptomatic infections transmit in our model. The first infection is symptomatic with probability *x*_0_. The chances of a host transmitting an infection declines exponentially with each infection at a rate *ξ* according to

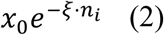

Epistatic interactions are modeled as a reduction in the infectivity of novel segment associations. Following a reassortment or introduction event the probability of infection is reduced by the following factor.

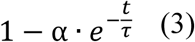

where t is the time since the reassortment or introduction event and α=0.5 and *τ* = 1/20 are the reduced compatibility magnitude and co-adaptation rate parameters.

Antigenically novel segments are introduced at a rate *i*_5_ in the context of an existing background present in the population. That is, an introduction brings into the population one new segment in the background of an existing constellation. These new segments are derived from an initial parent segment at the same loci sampled at the beginning of the simulation, to reflect their distant coalescence, possibly in a source animal or region.

Infected hosts are exposed to additional new infections. A co-infection may involve multiple circulating serotypes, and repeated infection with the same serotype. When a host infected with multiple serotypes transmits an infection, a single infecting serotype is generated and selected from the existing infecting serotypes. For the infecting serotype, a random parent serotype is selected, and with probability ρ each segment in this serotype is replaced by a random segment allele at the same loci, from all the infecting serotypes (including the parent strain).

A fraction (2 · 10^−4^) of infected hosts is sampled daily in the simulation, starting at the end of an initialization period lasting 50 years. When a host is sampled all infecting strains and segments are recorded. In addition, the immune history of a fraction (4 · 10^−6^) of all hosts (infected and non-infected) is sampled daily. Prevalence in simulated cumulative and frequency plots, is aggregated at two-year time windows and interpolated within windows using cubic interpolation. To capture epidemic dynamics, **Figure 1**A samples were aggregated at a shorter time period of six months.

### Statistical information

A two-tailed T-test was used to compare the association diversity of established and more recent G types (N=8). A likelihood ratio test comparing a single normal-distribution with two sharing a common variance was used when comparing the number of associations of different P types (N=3), with limitations at this sample size. The two comparisons (P-types and G-types) are not completely independent.

Segment overlap was calculated as the fraction of shared segments between pairs of randomly sampled serotypes when sampling with returns and discarding samples with identical serotype. The significance of overlap statistics (o) in individual countries and globally was determined by comparison to the same number of serotypes assembled from a random collection of G and P types sampled from the global pool. P-values which include a less than (<) sign, relate to limitations based on the number of randomizations (p<2/N) performed.

P-values for identifying G-P pair abundance being below or above random expectation were calculated using a two tailed Fisher’s exact test.

### Data Availability

The ‘strain IDs’ of rotavirus genomes used in the analysis are included in a supplementary excel file. Details about the selection criteria for sequences can be found in Woods et al.^27^.

### Code availability

Code for simulations is available at github.com/dzinder/segmentree.

## SUPPLEMENT

**Figure S1.**
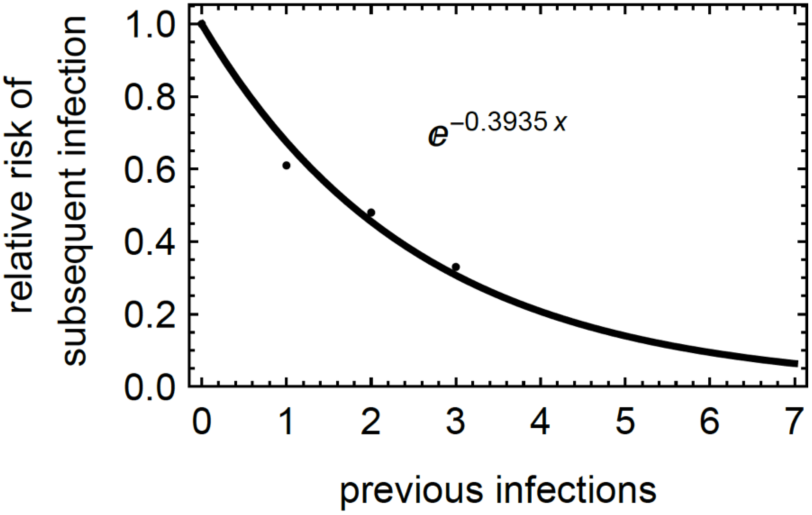
Relative risk of subsequent infection modeled as an exponential decline with previous infections. . Relative risk of subsequent infection in an Indian birth cohort was evaluated using data from Gladstone et al. 2011^43^ and fitted with an exponential curve. The rate (σ_gen_=0.3935) offers an upper value estimate for the effect of generalized immunity on infection risk in this cohort.

**Figure S2.**
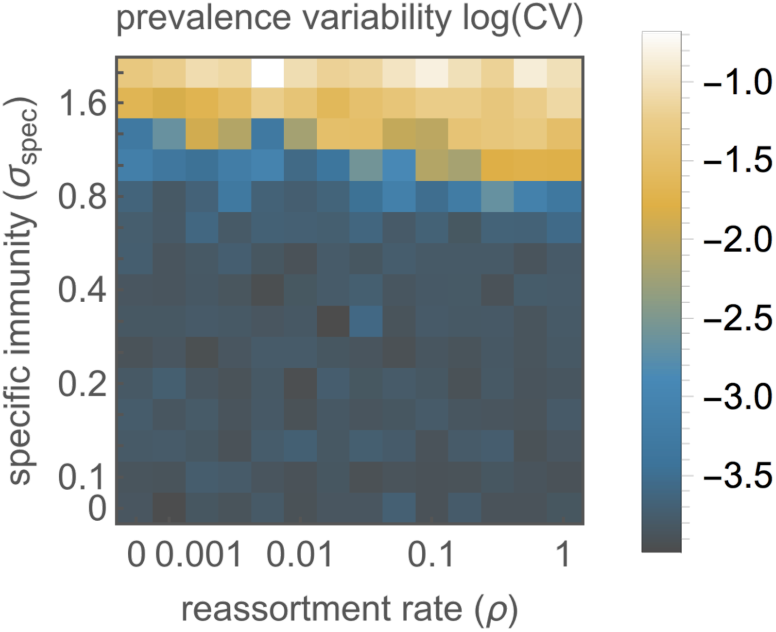
Prevalence variability with varying specific immunity and reassortment rates, where antigenic novelty is driven by introductions from an unlimited pool. Natural log of the prevalence coefficient of variation (CV) plotted as a function of the strength of specific immunity and of the reassortment rate. The CV was calculated as standard deviation divided by mean prevalence, when averaging two-year time intervals. Higher specific immunity causes increased prevalence variability and is associated with a regime which includes the replacement of strains rather than the coexistence of multiple strains. Reassortment increases incidence variability only for intermediate levels of specific immunity, with an effect on serotype diversity but not on incidence variability for lower levels of specific immunity. (parameters: β_gen_=3.51, σ_gen_=0.4, i_I_=8 year^-1^).

**Figure S3.**
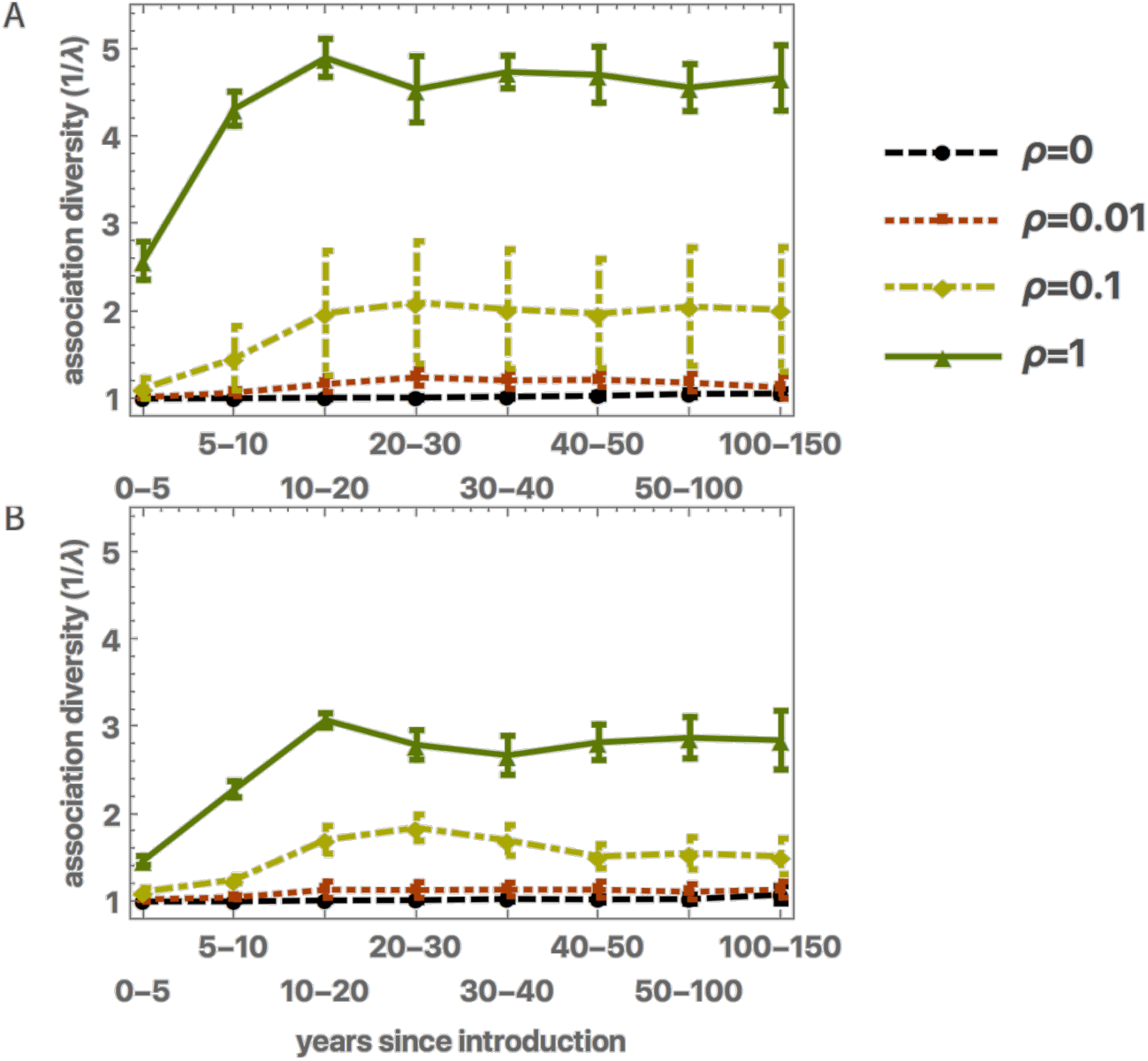
Segment associations following introduction with varying rates of reassortment. The diversity of associations (1/λ or ^2^D), with segments at different loci, a segment experiences following its introduction. Results were averaged across 6 independent runs for each data point. **A.** A newly introduced segment has a selective advantage which in the presence of reassortment can promote its association with more backgrounds. The number of associations a (non-extinct) newly introduced segment has reached an equilibrium level dependent on the reassortment rate **B** When considering epistatic interactions (see discussion) generating a fitness loss (50%) and regain (τ=1/20 year^-1^) following reassortment. (parameters: β = 3.51, σgen = 0.4, σspec = 0.25, iI=8 year^-1^)

**Figure S4.**
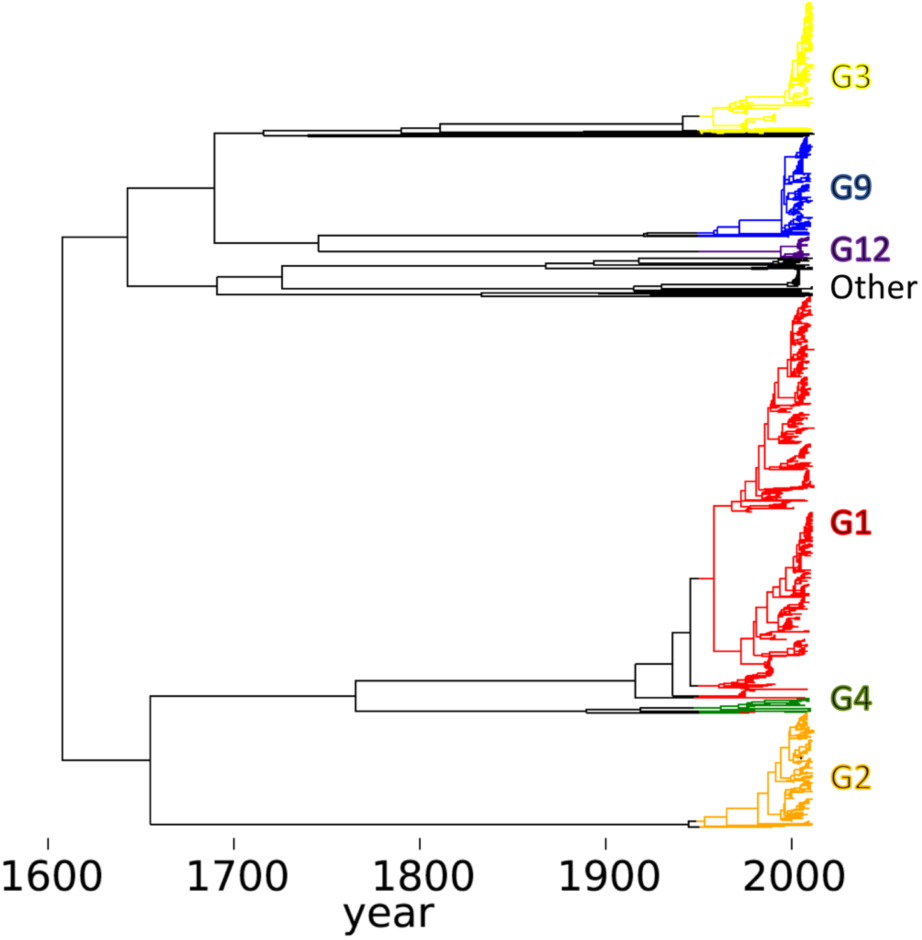
The phylogeny of Rotavirus VP7. The time-resolved phylogenetic tree of rotavirus A, VP7 segment (G protein) in humans. A single representative phylogeny of VP7 from the posterior distribution of trees, generated from a sample of 769 annotated GenBank sequence ^27^ using the phylogenetic inference package BEAST^62^, and color-coded by serotype.

**Figure S5.**
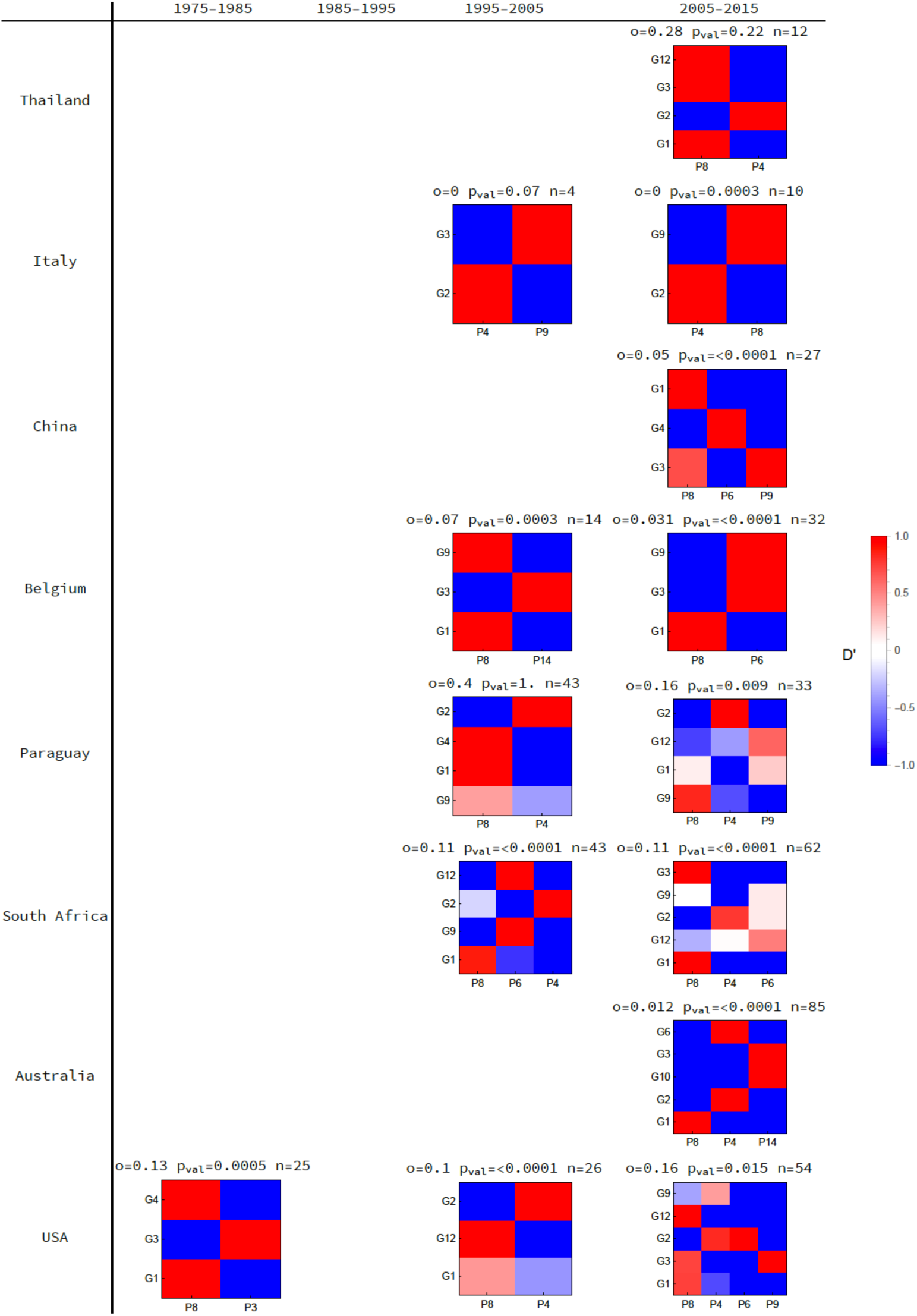
Deviation from equilibrium frequency and expected overlap for publically available G-P genotypes in different countries. The linkage disequilibrium (D’) between different G-P genotype pairs in samples from a specified country during decade long time intervals (1975-2015) was calculated using a curated sample of publically available sequence data (Woods et al. 2015). The overlap (o) of G-P genotypes sampled (n) in a country during a specified decade was calculated. The overlap represents the average fraction of shared segments between pairs of non-identical genotypes. Country level overlap was compared to the overlap in a random assembly of global segment alleles of the same sample size, the fraction of cases in which such an assembly resulted in a lower or equal overlap is reported (pval). P and G segments are ordered by their prevalence in each country and decade. Decades with at least two different genotypes were included. (Thailand, Italy, China, Belgium, Paraguay, South-Africa, USA).

**Figure S6.**
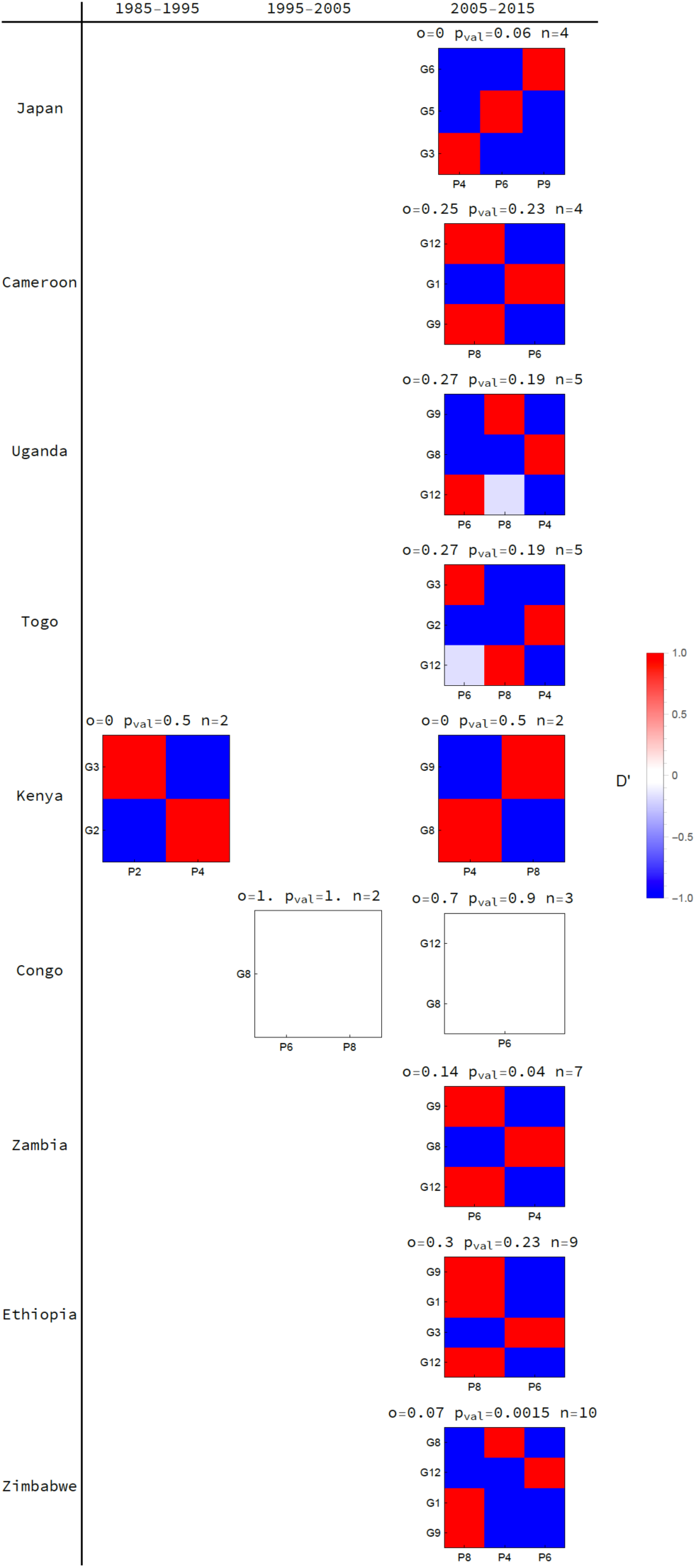
Deviation from equilibrium frequency and expected overlap for publically available G-P genotypes in different countries. The linkage disequilibrium (D’) between different G-P genotype pairs in samples from a specified country during decade long time intervals (1985-2015) was calculated using a curated sample of publically available sequence data (Woods et al. 2015). The overlap (o) of G-P genotypes sampled (n) in a country during a specified decade was calculated. The overlap represents the average fraction of shared segments between pairs of non-identical genotypes. Country level overlap was compared to the overlap in a random assembly of global segment alleles of the same sample size, the fraction of cases in which such an assembly resulted in a lower or equal overlap is reported (p_val_). P and G segments are ordered by their prevalence in each country and decade. Decades with at least two different genotypes were included. (Japan, Cameron, Uganda, Togo, Kenya, Congo, Zambia, Ethiopia, Zimbabwe).

**Table S1.** Frequency of G, P and G-P type combinations in curated publically available human sequence data^27^.

| G type | samples (%) | P type | samples (%) | G-P type | samples (%) |
| --- | --- | --- | --- | --- | --- |
| G1 | 340 (51.0%) | P[8] | 476 (71.5%) | G1P[8] | 329 (49.4%) |
| G2 | 101 (15.1%) | P[4] | 116 (17.4%) | G2P[4] | 97 (14.6%) |
| G3 | 72 (10.8%) | P[6] | 54 (8.1%) | G3P[8] | 55 (8.2%) |
| G9 | 66 (9.9%) | P[9] | 14 (2.1%) | G9P[8] | 50 (7.5%) |
| G12 | 39 (5.9%) | P[14] | 3 (0.5%) | G4P[8] | 22 (3.3%) |
| G4 | 25 (3.8%) | P[3] | 2 (0.3%) | G12P[8] | 18 (2.7%) |
| G8 | 16 (2.4%) | other | 2 (0.3%) | G12P[6] | 14 (2.1%) |
| G6 | 5 (0.8%) |  |  | G9P[6] | 13 (2.0%) |
| other | 3 (0.4%) |  |  | G8P[4] | 9 (1.4%) |
|  |  |  |  | G8P[6] | 6 (0.9%) |
|  |  |  |  | G1P[6] | 6 (0.9%) |
|  |  |  |  | G3P[6] | 6 (0.9%) |
|  |  |  |  | G12P[9] | 5 (0.8%) |
|  |  |  |  | G3P[9] | 4 (0.6%) |
|  |  |  |  | other | 33 (4.9%) |
| total |  |  |  |  | 667 (100%) |

## REFERENCES

1. Grenfell, B. T. Unifying the Epidemiological and Evolutionary Dynamics of Pathogens. Science 303, 327–332 (2004).

2. Gupta, S. et al. The maintenance of strain structure in populations of recombining infectious agents. Nat Med 2, 437–442 (1996).

3. Lipsitch, M. & O'Hagan, J. J. Patterns of antigenic diversity and the mechanisms that maintain them. J. R. Soc. Interface 4, 787–802 (2007).

4. Buckee, C. O., Recker, M., Watkins, E. R. & Gupta, S. Role of stochastic processes in maintaining discrete strain structure in antigenically diverse pathogen populations. Proc. Natl. Acad. Sci. 108, 15504–15509 (2011).

5. Artzy-Randrup, Y. et al. Population structuring of multi-copy, antigen-encoding genes in Plasmodium falciparum. eLife 1, (2012).

6. Zinder, D., Bedford, T., Gupta, S. & Pascual, M. The Roles of Competition and Mutation in Shaping Antigenic and Genetic Diversity in Influenza. PLoS Pathog 9, e1003104 (2013).

7. Cobey, S. Pathogen evolution and the immunological niche: Evolution of pathogen diversity. Ann. N. Y. Acad. Sci. 1320, 1–15 (2014).

8. Gause, G. F. The struggle for existence, 163 pp. Williams Wilkins Baltim. (1934).

9. Hutchinson, G. E. The multivariate niche. in Cold Spr. Harb. Symp. Quant. Biol 22, 415–421 (1957).

10. MacArthur, R. & Levins, R. The limiting similarity, convergence, and divergence of coexisting species. Am. Nat. 377–385 (1967).

11. Abrams P. *The theory of limiting similarity*. 14, (Annual Reviews, 1983).

12. Scheffer, M. & van Nes, E. H. Self-organized similarity, the evolutionary emergence of groups of similar species. Proc. Natl. Acad. Sci. 103, 6230–6235 (2006).

13. Cavender-Bares, J., Kozak, K. H., Fine, P. V. A. & Kembel, S. W. The merging of community ecology and phylogenetic biology. Ecol. Lett. 12, 693–715 (2009).

14. HilleRisLambers, J., Adler, P. B., Harpole, W. S., Levine, J. M. & Mayfield, M. M. Rethinking community assembly through the lens of coexistence theory. Annu. Rev. Ecol. Evol. Syst. 43, 227–248 (2012).

15. Chesson, P. Mechanisms of maintenance of species diversity. Annu. Rev. Ecol. Syst. 31, 343–366 (2000).

16. Kapikian, A. Z. A rotavirus vaccine for prevention of severe diarrhoea of infants and young children: development, utilization and withdrawal. in Novartis Foundation Symposium 153–179 (Chichester; New York; John Wiley; 1999, 2001).

17. Matthijnssens, J. et al. Full Genome-Based Classification of Rotaviruses Reveals a Common Origin between Human Wa-Like and Porcine Rotavirus Strains and Human DS-1-Like and Bovine Rotavirus Strains. J. Virol. 82, 3204–3219 (2008).

18. Ward, R. L., Clark, H. F. & Offit, P. a. Influence of potential protective mechanisms on the development of live rotavirus vaccines. J. Infect. Dis. 202 Suppl, S72–9 (2010).

19. Offit, P. a. Host factors associated with protection against rotavirus disease: the skies are clearing. J. Infect. Dis. 174 Suppl, S59–64 (1996).

20. Crawford, S. E. et al. Protective Effect of Natural Rotavirus Infection in an Indian Birth Cohort. 337–346 (2011).

21. Velazquez, F. R. et al. Cohort study of rotavirus serotype patterns in symptomatic and asymptomatic infections in Mexican children. Pediatr. Infect. Dis. J. 12, 54–61 (1993).

22. Leshem, E. et al. Distribution of rotavirus strains and strain-specific effectiveness of the rotavirus vaccine after its introduction: a systematic review and meta-analysis. Lancet Infect. Dis. 14, 847–856 (2014).

23. Matthijnssens, J. et al. Phylodynamic analyses of rotavirus genotypes G9 and G12 underscore their potential for swift global spread. Mol. Biol. Evol. 27, 2431–2436 (2010).

24. Arista, S. et al. Heterogeneity and Temporal Dynamics of Evolution of G1 Human Rotaviruses in a Settled Population. J. Virol. 80, 10724–10733 (2006).

25. McDonald, S. M., Davis, K., McAllen, J. K., Spiro, D. J. & Patton, J. T. Intra-genotypic diversity of archival G4P [8] human rotaviruses from Washington, DC. Infect. Genet. Evol. 11, 1586–1594 (2011).

26. McDonald, S. M. et al. Evolutionary Dynamics of Human Rotaviruses: Balancing Reassortment with Preferred Genome Constellations. PLoS Pathog. 5, e1000634 (2009).

27. Woods, R. J. Intrasegmental recombination does not contribute to the long-term evolution of group A rotavirus. Infect. Genet. Evol. 32, 354–360 (2015).

28. Santos, N. & Hoshino, Y. Global distribution of rotavirus serotypes/genotypes and its implication for the development and implementation of an effective rotavirus vaccine. Rev. Med. Virol. 15, 29–56 (2005).

29. Heiman, E. M. et al. Group A human rotavirus genomics: evidence that gene constellations are influenced by viral protein interactions. J. Virol. 82, 11106–11116 (2008).

30. Lipsitch, M., Colijn, C., Cohen, T., Hanage, W. P. & Fraser, C. No coexistence for free: neutral null models for multistrain pathogens. Epidemics 1, 2–13 (2009).

31. Lipsitch, M., Siller, S. & Nowak, M. A. The evolution of virulence in pathogens with vertical and horizontal transmission. Evolution 1729–1741 (1996).

32. Levin, S. & Pimentel, D. Selection of intermediate rates of increase in parasite-host systems. Am. Nat. 308–315 (1981).

33. van Veelen, M., Luo, S. & Simon, B. A simple model of group selection that cannot be analyzed with inclusive fitness. J. Theor. Biol. 360, 279–289 (2014).

34. Maslov, S. & Sneppen, K. Population cycles and species diversity in dynamic Kill-the-Winner model of microbial ecosystems. Sci. Rep. 7, (2017).

35. McDonald, S. M. et al. Diversity and relationships of cocirculating modern human rotaviruses revealed using large-scale comparative genomics. J. Virol. 86, 9148–62 (2012).

36. Afrad, M. et al. Changing profile of rotavirus genotypes in Bangladesh, 2006--2012. BMC Infect. Dis. 13, 320 (2013).

37. De Grazia, S. et al. Data mining from a 27-years rotavirus surveillance in Palermo, Italy. Infect. Genet. Evol. J. Mol. Epidemiol. Evol. Genet. Infect. Dis. 1–8 (2014). doi:10.1016/j.meegid.2014.03.001

38. Nakagomi, T. et al. Apparent extinction of non-G2 rotavirus strains from circulation in Recife, Brazil, after the introduction of rotavirus vaccine. Arch. Virol. 153, 591–593 (2008).

39. Kirkwood, C. D., Boniface, K., Barnes, G. L. & Bishop, R. F. Distribution of Rotavirus Genotypes After Introduction of Rotavirus Vaccines, Rotarix^®^ and RotaTeq^®^, into the National Immunization Program of Australia. Pediatr. Infect. Dis. J. 30, S48–S53 (2011).

40. Zinder, D., Woods, R. J. & Pascual, M. Early signs of post-vaccination change in the USA rotavirus population through mutation, migration and shifting prevalence. in (Ecological Society of America, 2014).

41. Bedford, T., Rambaut, A. & Pascual, M. Canalization of the evolutionary trajectory of the human influenza virus. BMC Biol. 10, 38 (2012).

42. Bedford, T. et al. Integrating influenza antigenic dynamics with molecular evolution. Elife 3, e01914 (2014).

43. Gladstone, B. P. et al. Protective effect of natural rotavirus infection in an Indian birth cohort. N. Engl. J. Med. 365, 337–346 (2011).

44. Cunliffe, N. A. et al. Molecular and Serologic Characterization of Novel Serotype G8 Human Rotavirus Strains Detected in Blantyre, Malawi. Virology 274, 309–320 (2000).

45. Cunliffe, N. A. et al. Rotavirus G and P types in children with acute diarrhea in Blantyre, Malawi, from 1997 to 1998: predominance of novel P [6] G8 strains. J. Med. Virol. 57, 308–312 (1999).

46. Rahman, M. et al. Predominance of rotavirus G9 genotype in children hospitalized for rotavirus gastroenteritis in Belgium during 1999-2003. J. Clin. Virol. 33, 1–6 (2005).

47. Yang, X.-L. et al. Temporal changes of rotavirus strain distribution in a city in the northwest of China, 1996-2005. Int. J. Infect. Dis. 12, e11–e17 (2008).

48. Rahman, M. et al. Evolutionary History and Global Spread of the Emerging G12 Human Rotaviruses. J. Virol. 81, 2382–2390 (2007).

49. Matthijnssens, J., Rahman, M., Ciarlet, M. & Van Ranst, M. Emerging human rotavirus genotypes. (2008).

50. Maes, P., Matthijnssens, J., Rahman, M. & Van Ranst, M. RotaC: a web-based tool for the complete genome classification of group A rotaviruses. BMC Microbiol. 9, 238 (2009).

51. Koelle, K., Cobey, S., Grenfell, B. & Pascual, M. Epochal Evolution Shapes the Phylodynamics of Interpandemic Influenza A (H3N2) in Humans. Science 314, 1898–1903 (2006).

52. Parrish, C. R., Murcia, P. R. & Holmes, E. C. Influenza Virus Reservoirs and Intermediate Hosts: Dogs, Horses, and New Possibilities for Influenza Virus Exposure of Humans: FIG 1. J. Virol. 89, 2990–2994 (2015).

53. Kryazhimskiy, S., Dushoff, J., Bazykin, G. A. & Plotkin, J. B. Prevalence of Epistasis in the Evolution of Influenza A Surface Proteins. PLoS Genet 7, e1001301 (2011).

54. Lindstrom, S. E., Cox, N. J. & Klimov, A. Evolutionary analysis of human H2N2 and early H3N2 influenza viruses: evidence for genetic divergence and multiple reassortment among H2N2 and/or H3N2 viruses. in International Congress Series 1263, 184–190 (Elsevier, 2004).

55. Webster, R. G. in Origin and evolution of viruses. 377–390 (Academic Press, San Diego, 1999).

56. Nachbagauer, R. et al. Defining the antibody cross-reactome directed against the influenza virus surface glycoproteins. Nat Immunol 18, 464–473 (2017).

57. Whitton, J. L., Cornell, C. T. & Feuer, R. Host and virus determinants of picornavirus pathogenesis and tropism. Nat. Rev. Microbiol. 3, 765–776 (2005).

58. Bourhy, H. et al. The origin and phylogeography of dog rabies virus. J. Gen. Virol. 89, 2673–2681 (2008).

59. Holmes, E. C. & Zhang, Y.-Z. The evolution and emergence of hantaviruses. Curr. Opin. Virol. 10, 27–33 (2015).

60. Pitzer, V. E. et al. Demographic Variability, Vaccination, and the Spatiotemporal Dynamics of Rotavirus Epidemics. Science 325, 290–294 (2009).

61. Velázquez, F. R. et al. Rotavirus Infection in Infants as Protection against Subsequent Infections. N. Engl. J. Med. 335, 1022–1028 (1996).

62. Drummond, A. & Rambaut, A. BEAST: Bayesian evolutionary analysis by sampling trees. BMC Evol. Biol. 7, 214 (2007).

